# Oxygen-releasing nanodroplets relieve intratumoral hypoxia in head and neck cancer spheroids

**DOI:** 10.1101/2024.05.26.595914

**Authors:** Marvin Xavierselvan, Brooke Bednarke, Ronak Tarun Shethia, Vicky Yang, Leah Moses, Srivalleesha Mallidi

**Author notes:** Authors have equal contribution (Brooke Bednarke, Ronak Tarun Shethia).

## Abstract

Hypoxia in solid tumors, including head and neck cancer (HNC), contributes to treatment resistance, aggressive phenotypes, and poor clinical outcomes. Perfluorocarbon nanodroplets have emerged as promising oxygen carriers to alleviate tumor hypoxia. However, a thorough characterization of the hypoxia alleviation effects in terms of sustenance of oxygenated environments have not been thoroughly studied. In this study, we developed and characterized perfluoropentane nanodroplets (PFP NDs) for co-delivery of oxygen and the photoactivatable drug or photosensitizer benzoporphyrin derivative (BPD) to hypoxic HNC spheroids. The PFP NDs exhibited excellent stability, efficient oxygen loading/release, and biocompatibility. Using 3D multicellular tumor spheroids of FaDu and SCC9 HNC cells, we demonstrated the ability of oxygenated PFP NDs to penetrate the hypoxic core and alleviate hypoxia, as evidenced by reduced fluorescence of a hypoxia-sensing reagent and downregulation of hypoxia-inducible factors HIF-1α and HIF-2α. BPD-loaded PFP NDs successfully delivered the photosensitizer into the spheroid core in a time-dependent manner. These findings highlight the potential of PFP NDs as a co-delivery platform to overcome hypoxia-mediated treatment resistance and improve therapy outcomes in HNC.

## 2. Introduction

Head and neck cancer (HNC) encompasses a heterogeneous group of malignancies originating from the mucosal epithelium of the head and neck regions, mainly in the oral cavity, including the lip, pharynx, larynx, nasal cavity, and paranasal sinuses.^1^ According to recent GLOBOCAN estimates, HNC accounts for roughly 900,000 new cases (3-5% of all cancer diagnoses) and roughly 450,000 deaths each year globally.^2^ The development and progression of HNC are closely linked to various risk factors, including tobacco and alcohol consumption, human papillomavirus infection, and genetic predisposition.^3,4^ At the molecular level, HNC exhibits substantial heterogeneity, with diverse genetic alterations and dysregulated signaling pathways contributing to its aggressive behavior and therapeutic resistance.^5-10^ Additionally, hypoxia, a hallmark of solid tumors,^11,12^ plays a pivotal role in shaping the tumor microenvironment (TME) and compromising treatment outcomes in HNC. Hypoxic conditions within tumors arise due to the rapid proliferation of cancer cells and inadequate supply of oxygen from the aberrant tumor vasculature.^13-15^ These hypoxic regions can lead to the stabilization and activation of hypoxia-inducible factors (HIFs), a family of transcription factors that regulate the expression of genes involved in angiogenesis, glycolysis, cell survival, and epithelial-mesenchymal transition (EMT).^16-19^ In HNC, elevated levels of HIF-1α have been associated with aggressive tumor phenotypes, treatment resistance, and poor clinical outcomes.^8^

The challenges posed by tumor hypoxia necessitate the development of more effective and targeted therapeutic approaches. Particularly, therapies such as radiotherapy and photodynamic therapy (PDT), that require oxygenated environments for effective outcomes to sustain the DNA damage and generate reactive oxygen species respectively.^20-23^ To alleviate hypoxia and improve treatment outcomes, various strategies ranging from micro and nanoformulations to hypoxia activated prodrugs have been employed.^24-28^ Amongst the nanomaterials that can specifically carry oxygen, perfluorocarbon (PFC)-based nanocarriers that act as artificial blood substitutes are attractive due to their high oxygen solubility and storage capacity, biocompatibility, and metabolic non-reactivity.^29,30^ Their primary use as oxygen carriers has been shown to crush hypoxia in solid tumors.^31-36^ Studies have shown the molecular ratio of dissolved oxygen in PFC is 40-56 mL O_2_ per 100 mL PFC at standard temperature and pressure.^27,37,38^ By encapsulating PFCs into nanoemulsions, nanodroplets can be designed to provide tunable, sustained oxygen release to boost tumor oxygenation, alleviate hypoxia, and improve PDT efficacy.^32-36,39,40^ Unlike hemoglobin’s localized chemical binding to oxygen molecules, PFC’s oxygen solubility is directly proportional to the gas’s partial pressure.^41^ Therefore, oxygen can be rapidly released from PFCs when needed based on concentration gradient or through triggered release such as acoustic droplet vaporization or light-triggered delivery.^42-44^ Recent studies have tested perfluorohexane nanodroplets (NDs) loaded with drugs such as photosensitizers (PSs) for enhanced photodynamic therapy.^34,45^ The oxygen carried by the NDs helped overcome hypoxia and enhance photodynamic effects.

PFC NDs ability to carry oxygen to tumors has previously been demonstrated in several *in vivo* models.^32,34,35,39^ However, thorough characterization of the hypoxia alleviation effects in terms of sustenance post NDs delivery are not thoroughly studied. In order to investigate the long-term effects and sustained alleviation of hypoxia by PFC NDs, physiologically relevant models that accurately recapitulate the tumor microenvironment are needed. Conventional 2D monolayer models do not provide the complexity of tumor heterogeneity and architectures, while animal models can resemble actual tumor environments but lack high throughput and are difficult to control.^6,46,47^ Three dimensional multicellular tumor spheroid models are a promising alternative that bridges the gap between traditional 2D monolayer cultures and *in vivo* animal models. Spheroids are self-assembled, multicellular aggregates grown in suspension that mimic tumor heterogeneity and complex architectures.^48-50^ Additionally, spheroid models recapitulate key features of solid tumors, such as cell-cell interactions, extracellular matrix production, nutrient and drug penetration gradients, and hypoxic regions. The presence of hypoxic regions within the spheroids allows for the evaluation of nanocarriers uptake and oxygen delivery in oxygen-deprived areas, which are associated with therapeutic resistance and metastatic potential.^50^ Through comprehensive characterization and evaluation in HNC spheroid models, this study seeks to elucidate the ability of the developed perfluoropentane (PFP) NDs to deliver oxygen and drugs and alleviate tumor hypoxia while also evaluating the drug delivery capability of NDs. By addressing the critical challenges posed by hypoxia and limited drug delivery, the developed nanodroplets system holds the potential to overcome treatment resistance and improve survival rates for patients with hypoxic and aggressive HNC tumors.

## 3. Materials and Methods

### 3.1. Synthesis of the Nanodroplets

The PFP NDs were synthesized using the water emulsion method. A combination of 1,2-dipalmitoyl-sn-glycero-3-phosphocholine (DPPC), 1,2-distearoyl-sn-glycero-3-phosphoethanolamine-N-[methoxy(polyethylene glycol)-2000] (DSPE-mPEG), 1,2-distearoyl-sn-glycero-3-phosphoethanolamine-N-[maleimide(polyethylene glycol)-2000] (DSPE-PEG(2000)-mal), and cholesterol were added to a 10 mL round-bottom flask at a molar ratio of 94.7:0.4:3:1.9, respectively. The chloroform was then evaporated using the nitrogen flow and rotary evaporator to create a dry film of lipids. The film was hydrated with deionized ultra-pure water to yield a final lipid concentration of 10 mg/mL. To this solution, 250 µL of n-perfluoropentane (Exfluor) was added and sonicated in a bath sonicator for 10 seconds at 10 °C, and vortexed for 10 seconds. This procedure was repeated until no visible PFP was observed or 10 minutes had passed. Finally, thiol-functionalized methoxy polyethylene glycol (mPEG-SH; M.W.: 30k) was added to the solution and mixed at 45 RPM overnight at 4 °C. The solution was then centrifuged at 250 RCF, and the supernatant was collected and stored at 4 °C as the final nanodroplets’ solution. For synthesizing PFP NDs loaded with benzoporphyrin derivative (BPD-PFP NDs), 0.45 mg of BPD in chloroform was incorporated into the lipid mixture and synthesized using the abovementioned procedure.

### 3.2. Stability of the Nanodroplets

The stability of the NDs were evaluated at 4 °C over 8 weeks. The hydrodynamic diameter, polydispersity indices (PDI), and zeta potential of the NDs were measured weekly in triplicate to assess any changes using a dynamic light scattering (DLS) system (NanoBrook ZetaPALS, Brookhaven Instruments). The NDs were diluted 1200x and 200x for hydrodynamic diameter and zeta potential measurements respectively. To investigate the potential effects of oxygen purging on the NDs size, DLS measurements were made before, immediately after, and 24 hours after being purged with oxygen (2 minutes at 0.5 lpm). All stability assessments were investigated using at least three independent batches of the NDs.

### 3.3. Optical Characterization of the Nanodroplets

A UV-visible spectrophotometer (Evolution 300, Thermo Fisher Scientific) was used to measure the concentration of BPD in the NDs. Both free BPD and BPD-PFP NDs were dissolved in dimethyl sulfoxide (DMSO) for UV-visible absorption spectrophotometry. Absorbance and fluorescence characterization were performed to evaluate the optical properties of the BPD-PFP NDs compared to free BPD in a quenched (Dulbecco’s phosphate buffered saline; PBS) and unquenched setting (DMSO). All the samples’ absorbance and fluorescence emission spectra were collected using Varioskan LUX microplate reader (Thermo Fisher Scientific). For fluorescence studies, the samples were excited with an excitation wavelength of 405 nm.

### 3.4. Measurement of Oxygen Loading of the Nanodroplets

For oxygen loading measurements, 0.65 mL of PFP NDs or PBS were purged with oxygen at a flow rate of 0.5 lpm for 2 minutes and then sealed to maintain the oxygen saturation. Then, 0.5 mL of the above-prepared solution was introduced into 4.5 mL of deoxygenated PBS pretreated with nitrogen (purged for 30 minutes), and a dissolved oxygen meter (FiveGo DO meter F4, Mettler Toledo) was used to record the oxygen content as a function of time.

### 3.5. Cell Culture

Human HNC cell lines FaDu (hypopharyngeal squamous cell carcinoma) and SCC9 (tongue squamous cell carcinoma) were obtained from the American Type Culture Collection and cultured in Dulbecco’s Modified Eagle Medium (DMEM) and was cultured in DMEM/F-12 media supplemented with 10% fetal bovine serum (Gibco) and 1% antibiotics (Penicillin and Streptomycin 1:1 v/v; Corning) respectively. The DMEM/F-12 media was also supplemented with 0.4 µg/mL hydrocortisone (Sigma-Aldrich). The cultured cells were tested for mycoplasma contamination and maintained at 37 °C and 5% CO_2_. The cells were sub-cultured and used for experiments while they were in the exponential growth phase.

### 3.6. Cellular Uptake of the Nanodroplets

For evaluating the cellular uptake of the PFP NDs, FaDu and SCC9 cells were seeded in a 24-well plate at a density of 50,000 cells per well and incubated with free BPD or BPD-PFP NDs at a concentration of 0.25 µM of BPD equivalent for 0, 1, 2, 3, 4, 6, 8, and 24 hours. After the incubation, cells were washed with ice-cold PBS twice and lysed using 0.2% Triton X-100 at 4°C. BPD fluorescence was measured using a microplate reader (Varioskan LUX; Thermo Fisher Scientific) with appropriate standards facilitating fluorescence intensity to be converted to moles of BPD. Protein concentration was quantified using the bicinchoninic acid (BCA) protein assay kit (Thermo Fisher Scientific) according to the manufacturer’s instructions. To verify the nanodroplet’s internalization using fluorescence imaging, cells were seeded in a 96-well black-walled plate at a density of 5,000 cells per well and incubated with Bodipy-tagged PFP NDs (1:100 dilution) for various time points. After incubation, cells were washed with PBS, fixed in formalin, stained with Hoechst (33342; Thermo Fisher Scientific), and imaged with the EVOS M7000 imaging system (Invitrogen) using a 40x objective. Hoechst and Bodipy fluorescence were imaged with a DAPI and GFP LED light cube. The fluorescence images were processed using ImageJ software (National Institutes of Health).

### 3.7. Cytotoxicity of the Nanodroplets

To evaluate the PFP NDs cytotoxicity (no light), FaDu and SCC9 cells were seeded in a 24-well plate at a density of 50,000 cells per well. The following day, cells were incubated with varying concentrations of BPD-PFP NDs for 1 hour or 0.25 µM of BPD-PFP NDs for various time points. After the respective time of incubation, cells were washed with PBS, and fresh media was added and incubated for a further 24 hours before performing the viability assay using 3-(4,5-dimethylthiazol-2-yl)-2,5-diphenyltetrazolium bromide (MTT, Thermo Fisher Scientific). The media was replaced with fresh cell culture media containing MTT (0.25 mg/mL) and incubated for 1 hour at 37 °C. The formazan crystals formed inside the cells were dissolved using DMSO, and the absorbance was recorded at 570 nm using a microplate reader (Varioskan LUX; Thermo Fisher Scientific). Cell viability was calculated as a percentage of absorbance at 570 nm compared to untreated controls.

### 3.8. Establishment of 3D Spheroids

To establish the multicellular tumor spheroids, FaDu and SCC9 cells were seeded in 96-well ultra-low attachment (ULA) U-bottom plates. To determine the optimal seeding density for further experiments, cells were seeded in ULA plates at various densities ranging from 1000 to 25,000 cells/well. The spheroids were monitored daily and imaged using a 4x objective and bright field on an EVOS M7000 imaging system.

### 3.9. Evaluation of Hypoxia Heterogeneity in 3D Spheroids

For evaluating the progression of hypoxia in the spheroids, ‘*in well’* staining method was used. FaDu spheroids on days 7, 10, 14, 17, and 21 were incubated with 100 µM of pimonidazole for one hour. After the incubation, the media was washed, and spheroids were fixed in 10% formalin for one hour. Once the fixation was completed, formalin was washed off with PBS, and the spheroids were permeabilized with 2% triton X-100 for one hour. After permeabilization, spheroids were washed three times with PBS. Then, the spheroids were incubated with antibody against pimonidazole adducts (1:50 dilution, Cat: HP7-100, Hypoxyprobe Red549 Kit) overnight at 4 °C. Later, the spheroids were washed off the excess antibody, and they were imaged with EVOS M7000 imaging system using a 4x objective and GFP LED light cube.

### 3.10. Hypoxia Alleviation in 3D Spheroids

For evaluating the hypoxia alleviation in the spheroids with the delivery of oxygenated PFP NDs, the spheroids were incubated with Image-iT™ Red Hypoxia Reagent overnight at a concentration of 5 µM. The next day, the spheroids were imaged for the hypoxia reagent fluorescence signals to obtain baseline measurements with the EVOS M7000 imaging system using a 4x objective and RFP LED light cube. After imaging, PBS and oxygenated PFP NDs were added to the spheroids at the dilution of 1:100 and incubated for 3 hours. After the time point, the spheroids were imaged for the hypoxia reagent fluorescence signals, and both the time point images were quantified in MATLAB with a custom-written script. The reduction in hypoxia was calculated as a difference in hypoxia reagent fluorescence between pre- and post-administration of the oxygenated PFP NDs.

### 3.11. Western Blotting

For validating the hypoxia alleviation in the spheroids western blotting was utilized. A total of 48 spheroids were collected in a 15 mL falcon tube for every treatment group. The spheroids were then centrifuged to separate out the media and lysed with 250 µL of NP-40 cell lysis buffer mixed with 1x protease and phosphatase inhibitor at 4 °C. The protein lysates were quantified with BCA assay. Equal amounts of protein (20 µg/well) were loaded on 10-20% Tris-Glycine gels (Invitrogen) and separated using sodium dodecyl sulfate-polyacrylamide gel electrophoresis. Proteins were transferred to a polyvinylidene fluoride membrane using an iBlot 2 gel transfer device (Invitrogen). The blots were later blocked in 5% non-fat dry milk in tris-buffered saline with 0.05% tween-20 for 1 hour and incubated with primary antibody against HIF-1α and HIF-2α overnight at 4 °C on a shaker. Proteins were detected using horseradish peroxidase-conjugated secondary antibody. The protein bands were developed using enhanced chemiluminescence detection reagent (Thermo Fisher Scientific) and imaged using a ChemiDoc MP imaging system (BioRad). Densitometric analysis of the blots was performed using ImageJ software to quantify the relative expressions of HIF-1α and HIF-2α.

### 3.12. Drug Delivery by the Nanodroplets

For studying the drug delivery by the NDs and the penetration into the core of the spheroids, free BPD and BPD-PFP NDs at a dilution of 0.5 µM (BPD eq. concentration) were added to spheroids and incubated for 1, 3, 6, and 24 hours. After the corresponding incubation period was completed, spheroids were washed off in PBS to remove excess BPD from the media. The spheroids were then imaged with the EVOS M7000 imaging system using a 4x objective and Qdot 705 LED light cube. Throughout the acquisition and processing of all the images, brightness and contrast levels were kept at the same level.

### 3.13. Primary Antibodies

Primary antibodies used in the study include: GAPDH (dilution: 1:1000, Cat: 97166, Cell Signaling Technology), β-Actin (dilution: 1:1000, Cat: 4970, Cell Signaling Technology), HIF-1α (dilution: 1:1000, Cat: 3716, Cell Signaling Technology), and HIF-2α (dilution: 1:1000, Cat: 7096, Cell Signaling Technology).

### 3.14. Statistical Analysis

Data analysis was performed using GraphPad Prism 10. The specific statistical tests used on data were mentioned in the figure captions. A *p*-value of less than 0.05 was considered statistically significant.

## 4. Results and Discussion

### 4.1. Characterization of Nanodroplets

Stable nanoparticles maintain their intended physicochemical properties over time, ensuring consistent performance and safety.^51,52^ In contrast, unstable nanoparticles can aggregate, fuse, or degrade, which can lead to changes in size, surface charge, and overall behavior. These alterations can compromise the therapeutic efficacy of the nanoparticles and may result in unintended toxicity, posing significant risks in clinical applications.^53^ Hence, we tested the stability of the fabricated NDs over the course of 8 weeks when stored at 4 °C (Fig. 1A). The NDs fabricated either as plain vehicles (PFP NDs, green line) for oxygen delivery or BPD-PFP NDs (Fig. 1A, blue line) for drug and oxygen co-delivery had an average hydrodynamic diameter and standard deviation of 215 nm ± 12 nm and 208 nm ± 18 nm, respectively, measured immediately after the synthesis. The NDs, irrespective of the type of encapsulation, showed an insignificant change in the hydrodynamic diameter throughout the duration of the stability study. These results agreed with our previously published report and other reports in the literature for similar types of PFC nanocarriers.^32,54,55^ The average PDI and standard deviation of the prepared PFP NDs and BPD-PFP NDs was 0.22 ± 0.03 and 0.24 ± 0.04, respectively (Fig. 1A). Insignificant differences in PDI were observed over the course of the stability study. The average zeta potential and standard deviation of the PFP NDs and BPD-PFP NDs on the day of preparation was -26.6 mV ± 1.0 mV and -29.4 mV ± 1.9 mV respectively (Fig. 1B). The surface charge of the NDs was negative due to most of the lipid composition being DPPC. The lipid DPPC is zwitterionic, causing it to expose its phosphate groups and create a negatively charged surface when it is in low ionic strength solutions like PBS and water.^56,57^ In addition to this, incorporating BPD in the NDs synthesis made the zeta potential even more negative, which was expected given that BPD nanoformulations are generally anionic in nature.^58,59^

**Figure 1:**
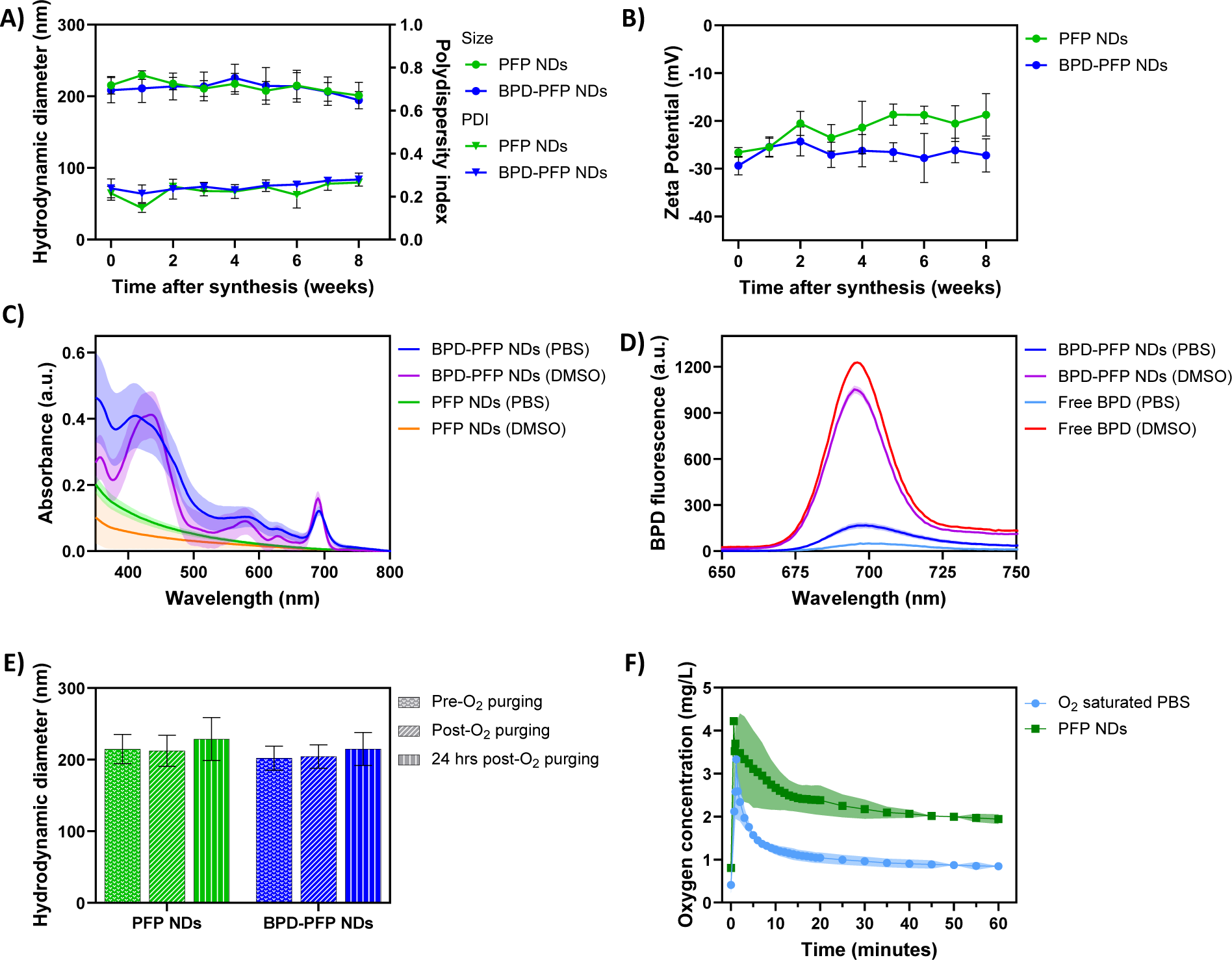
Physiochemical characterization of PFP NDs (plain, no-drug NDs) and BPD-PFP NDs (drug-loaded NDs). A) Hydrodynamic diameter and polydispersity indices and B) zeta potential of PFP NDs and BPD-PFP NDs monitored over 8 weeks post-synthesis. C) Absorbance spectra of PFP NDs and BPD-PFP NDs in PBS and DMSO. D) Fluorescence spectra of BPD-PFP NDs and free BPD in quenched (PBS) and unquenched (DMSO) settings. E) Hydrodynamic diameter of PFP NDs and BPD-PFP NDs before and after purging with oxygen. F) Dissolved oxygen concentration profile as a function of time for oxygen-saturated PBS and PFP NDs. For A-F), the results are expressed as mean ± standard deviation.

The optical properties (absorbance and fluorescence) of the synthesized NDs are shown in Figs. 1C and 1D, respectively. The absorbance curves for the PFP NDs in PBS and DMSO are shown in Fig. 1C by the green and orange lines, which highlight that the NDs without any BPD exhibit an exponentially decreasing absorbance from lower to higher wavelengths, which is consistent with previously reported optical properties of lipids.^60,61^ For NDs encapsulated with drug BPD, the Soret band (near 420 nm) and Q band (around 690 nm) of BPD was clearly visible in the absorbance spectrum. When the NDs were dissolved in DMSO, the lipid shell gets disrupted, allowing the Soret band of BPD to be more distinct, as shown in Fig.1C (purple line) near the 420 nm wavelength range.

Several previous studies have shown that porphyin drugs such as BPD when encapsulated in a nanoformulation have maintained their photoactivity but could be prone to self-quenching depending on the nanopackaging at high concentrations.^58^ To further understand if the drug BPD when encapsulated within the NDs was quenched, we performed fluorescence of them in comparison to free BPD (Fig. 1D). Since BPD is a hydrophobic molecule, free BPD aggregates and quenches in PBS, reducing fluorescence emission (light blue line, Fig. 1D). When dissolved in DMSO, free BPD completely de-quenches, and we can observe the restored fluorescence spectrum (red line, Fig. 1D). The effects of PBS on free BPD resulted in 96.7% ± 0.72% quenching of the peak fluorescence emission at 690 nm or only a 3.3% photoactivity as expected. Alternatively, the BPD-PFP NDs demonstrated only 85.1% ± 3.85% fluorescence quenching at 690 nm compared to its DMSO dispersion, or in other words 14.9% photoactivity (fluorescence emission in PBS/ fluorescence emission in DMSO). These results were encouraging as our BPD-PFP-NDs had photoactivity similar to the clinical liposomal formulation for BPD (Visudyne) at 8.83%. These results agreed with other findings of this porphyrin for both free and nanoformulations, supporting that encapsulating PS in NDs helps partially preserve their photoactivity.^58,62^

PFCs are known to have high oxygen-carrying capacity due to their high electronegativity of fluorine atoms.^41^ Therefore, when PFCs are encapsulated in the nanocarriers, this property can be exploited for oxygen delivery. It is important to characterize the NDs for their oxygen loading and release capacity, along with changes in size when encapsulated with oxygen. We show the stability of both PFP NDs and BPD-PFP NDs in terms of their hydrodynamic diameter after purging with oxygen (Fig. 1E green and blue bars, respectively). The DLS measurements revealed that there were no statistically significant changes in size immediately after oxygen purging and 24 hours post-purging. This was consistent for both PFP NDs and BPD-PFP NDs, and the results were congruent with other reported studies.^63,64^ After establishing the stability with respect to oxygen purging, we characterized the oxygen loading capacity of PFP NDs. From Fig. 1F, we can observe that the PFP NDs, after purging, can carry more oxygen than the oxygen-saturated PBS. Oxygenated PFP NDs or PBS were added to deoxygenated PBS and monitored with a dissolved oxygen meter. The dissolved oxygen concentration increased rapidly, as expected after the addition of NDs, and showed a gradual release of oxygen over time before reaching an equilibrium. PFP NDs showed a dissolved oxygen concentration of 1.94 mg/L compared to 0.84 mg/L for oxygen-saturated PBS. This observed trend was consistent with the other published reports.^65,66^ Based on these characterizations, we can conclude that the synthesized NDs were stable, retained photoactivity similar to clinical formulation of the drug, could be loaded with oxygen without change in size for up to 24 hours, and were stable when stored at 4 °C.

### 4.2. Cellular Characterization of the Nanodroplets

The cytotoxicity and biocompatibility of the NDs were assessed in FaDu and SCC9 cells using the MTT assay (Fig. 2A&B). BPD-PFP NDs had exhibited no toxicity in both the cell lines when incubated 1 hour with 10 µM BPD equivalent NDs. No statistically significant difference in cell viability was observed at all concentrations evaluated (Fig. 2A). Based on the above data and in congruence with previously reported literature,^32,67,68^ a BPD concentration of 0.25 µM was chosen for further studies. We incubated BPD-PFP NDs at the concentration of 0.25 µM BPD for varying time points ranging from 1 to 24 hours. BPD-PFP NDs displayed no inhibitory effect on the growth of the cells (Fig. 2B). These results show that the NDs are nontoxic and biocompatible and agree with previously reported studies.^69,70^

**Figure 2:**
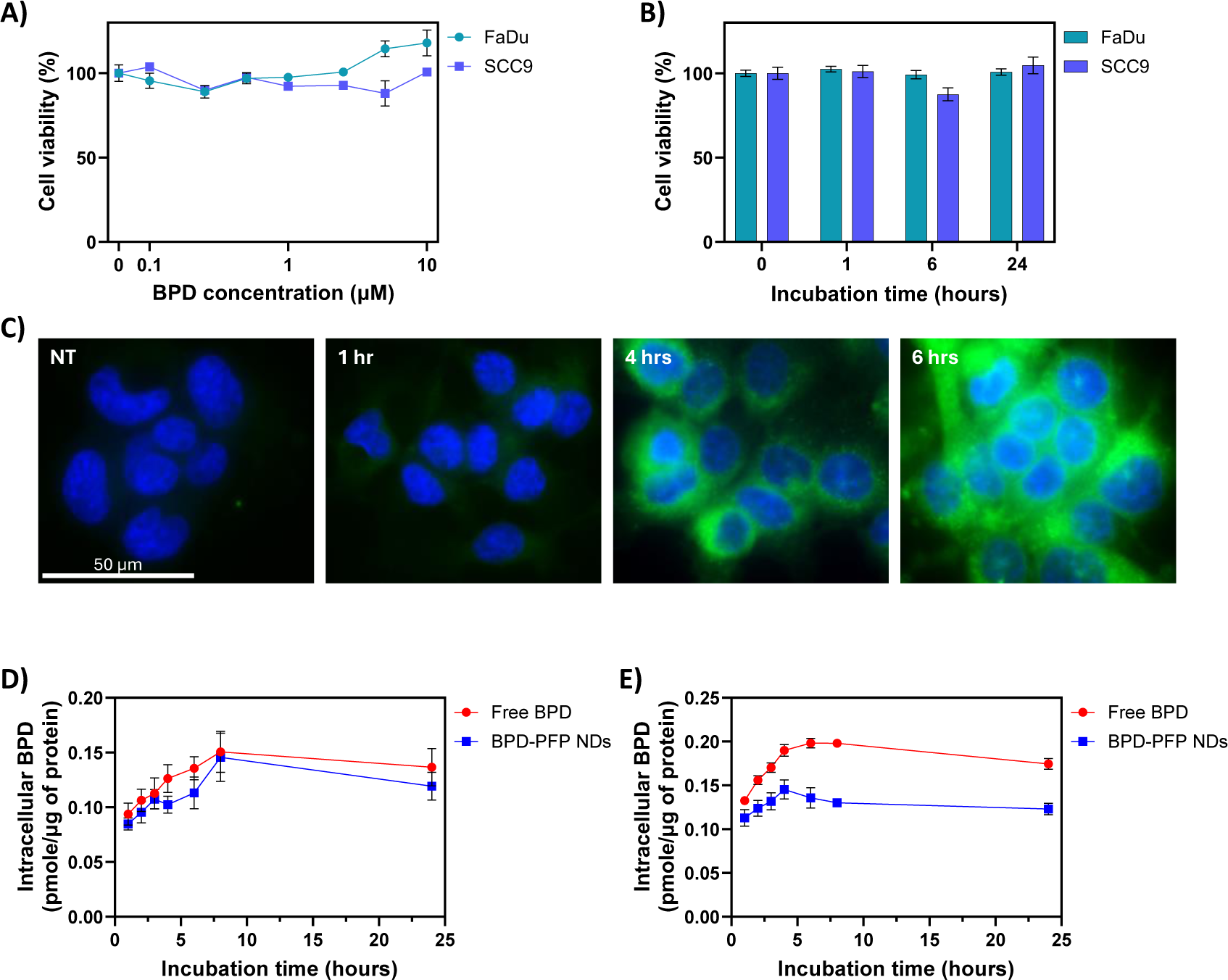
Evaluation of the cytotoxicity and cellular uptake of NDs over time. A) Concentration-dependent toxicity of BPD-PFP NDs after 1-hour incubation (n = 3). B) Time-dependent toxicity profile of BPD-PFP NDs at 0.25 µM concentration (n = 3). C) Time-lapse microscopy images showing the internalization of Bodipy-tagged PFP NDs in FaDu cells. (Scale bar = 50 µm). D&E) Quantitative analysis of free BPD or BPD-PFP NDs uptake via extraction assay in FaDu and SCC9 cells (n = 4). For A-B, D-E) the results are expressed as mean ± standard error.

Next, we investigated the cellular internalization of NDs (Fig 2C). We modified the plain PFP NDs by substituting a part of cholesterol with BODIPY-tagged cholesterol while forming the lipid film to evaluate the uptake of NDs in cells. From Fig. 2C, we can clearly observe that a time dependent increase in BODIPY fluorescence signal indicative of increased accumulation of NDs in FaDu cells. Further, to demonstrate that NDs are also delivering drugs intracellularly, we investigated the delivery of BPD by the PFP NDs in both FaDu and SCC9 cell lines (Fig. 2D&E) and compared the results with free BPD. The total intracellular BPD concentration is reported in pmol/µg of protein. In FaDu cells, both free BPD and BPD-PFP NDs exhibited similar uptake profiles, with increasing intracellular concentrations over time, peaking at the 8-hour time point. This suggests that the PFP NDs effectively delivered BPD into FaDu cells, achieving comparable uptake to free BPD. However, in SCC9 cells, the uptake of BPD-PFP NDs was lower than that of free BPD. The intracellular concentration of BPD delivered by PFP NDs peaked around the 4-hour time point, while free BPD reached its maximum uptake at approximately 6 hours. The lower uptake of BPD-PFP NDs in SCC9 cells could be attributed to the stealth function of PEG^71^ or the highly negative zeta potential of the NDs, which may influence their interaction with the cell membrane and subsequent internalization. Interestingly, SCC9 cells demonstrated higher uptake of both free BPD and BPD-PFP NDs compared to FaDu cells. This difference in uptake between the two cell lines could be due to variations in the expression levels of transporters, endocytic pathways, or intracellular processing mechanisms. Further investigation into the specific cellular drug transport pathways in these cell lines is warranted to better understand the observed differences in uptake.

### 4.3. Generation of Uniform-sized HNC Spheroid and Growth Characterization

HNC spheroids were generated using ULA plates, with each well initially producing a single spheroid of consistent size. The ULA plates resisted cell adhesion and promoted cell interaction, leading to aggregation and formation of a single spheroid per well.^72^ The ULA plates facilitated a highly controlled environment, resulting in spontaneous spheroid formation for FaDu and SCC9 cells in less than a day. This method proved to be more reproducible and reliable than other techniques for producing HNC tumor spheroids, yielding uniform spheroid size at the start of culture. Figure 3A shows representative FaDu and SCC9 spheroids imaged with a 4x objective using the EVOS M7000 imaging system. The cells in the spheroids grow in multiple layers, forming aggregates and leading to the formation of zones, namely an outer proliferating zone, a middle quiescent viable zone, and an inner hypoxic to necrotic core.^6,72^ The formation of different zones results in differential distribution of oxygen, nutrients, and growth factors and ultimately leads to heterogeneous uptake of drugs and drug resistance.

**Figure 3:**
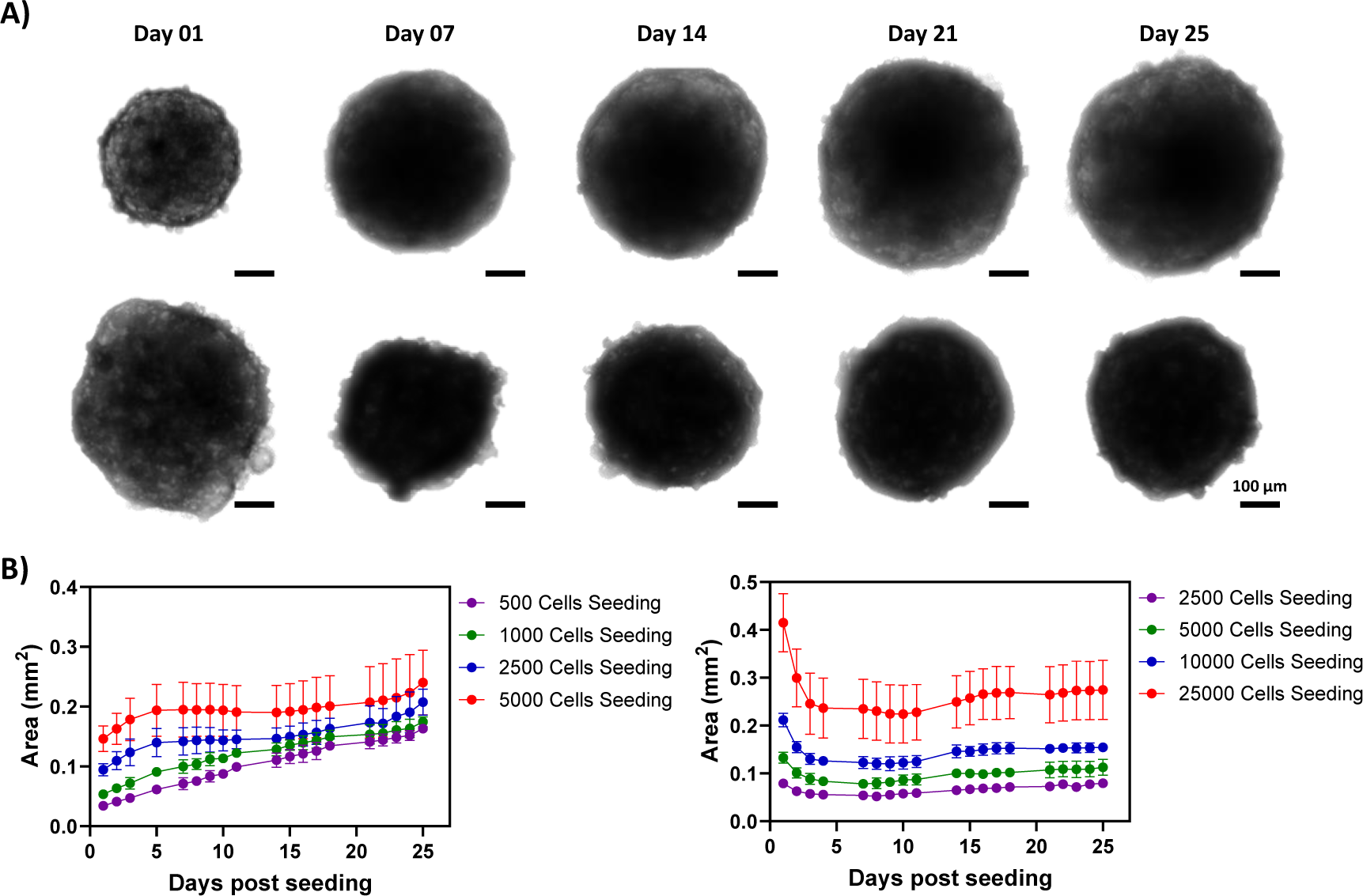
Characterization of HNC 3D spheroids. A) Bright field microscopy images showing the morphological changes of FaDu and SCC9 spheroids over time. (Scale bar = 100 µm). B) Growth kinetics of FaDu and SCC9 spheroids cultured from different initial seeding densities, demonstrating concentration-dependent growth profiles (n = 24). For B), the results are expressed as mean ± standard deviation.

To optimize seeding density for further experiments, FaDu and SCC9 cells were seeded at different initial cell seeding concentrations and cultured for 25 days. The spheroid diameter was calculated from the bright field images analyzed using a customized MATLAB script. The initial diameter of the spheroid depended on the cell seeding concentration, with the higher seeding group having the larger diameter. For FaDu spheroids, within each cell seeding group, the spheroids were uniform in size during the first week of culture (Fig. 3B). This uniformity could be attributed to the seeded cells aggregating together to form the spheroids on day 0. After the second week, the spheroids’ border became irregular and disorganized, suggesting cell proliferation in the outer layer. This led to larger spheroids with high variability in diameter by weeks three and four across all seeding groups despite the initial uniform sizes within each group. The random proliferation substantially increased spheroid size over time beyond the second week (Fig. 3B). In contrast, SCC9 spheroids reached their highest diameter on day 1. They progressively tightened their core in the first week of culture, resulting in a reduction in the diameter, and stabilized thereafter (Fig. 3C). Based on the results for subsequent investigations, HNC spheroids were produced with 2500 and 10000 cells for FaDu and SCC9 respectively because at these densities the spheroid diameters for both the cell lines were similar. The differences in growth patterns between FaDu and SCC9 spheroids highlight the importance of optimizing the culture conditions for each cell line. FaDu spheroids exhibited a more pronounced proliferative phase, leading to an increase in size and variability over time, while SCC9 spheroids reached a stable size more quickly. These findings underscore the need to consider cell line-specific characteristics, particularly with evaluating alleviation of hypoxia by NDs.

### 4.4. Time Course of Hypoxic Core Formation in FaDu Spheroids

Tumor hypoxia is a well-recognized phenomenon that contributes to treatment resistance, aggressive phenotypes, and poor clinical outcomes.^17,19^ In spheroid models, which closely mimic the 3D architecture and microenvironment of solid tumors, the onset and progression of hypoxia is closely related to spheroid size, as the diffusion of oxygen becomes limited beyond a certain radius (typically >200-500 µm).^50,73,74^ Monitoring the temporal progression of hypoxia is crucial, as it provides valuable insights into the dynamics of the tumor microenvironment and informs the optimal timing for therapeutic interventions targeting hypoxic regions.

To assess the progression of hypoxic core formation in FaDu spheroids, we performed ‘in well’ staining for pimonidazole adducts, a widely used marker for detecting tumor hypoxia. The IF/bright field overlay images in Fig. 4A depict the development of hypoxia over time, as indicated by the presence of pimonidazole signals. Consistent with previously reported works on other spheroids models,^74,75^ hypoxia began to form around day 10 or when the diameter reached around 500 µm, with the core becoming abundantly hypoxic approximately two weeks after spheroid formation. After two weeks, the hypoxic core remained at a similar level on subsequent measurement days, suggesting a stable hypoxic microenvironment once established.^76^ Quantification of pimonidazole-positive pixels using a MATLAB script revealed a statistically significant 40% hypoxic area on day 14 compared to the control (Fig. 4B) Based on these results, we selected day 14 as the optimal timepoint to treat the HNC spheroids with NDs and assess their performance in alleviating hypoxia.

**Figure 4:**
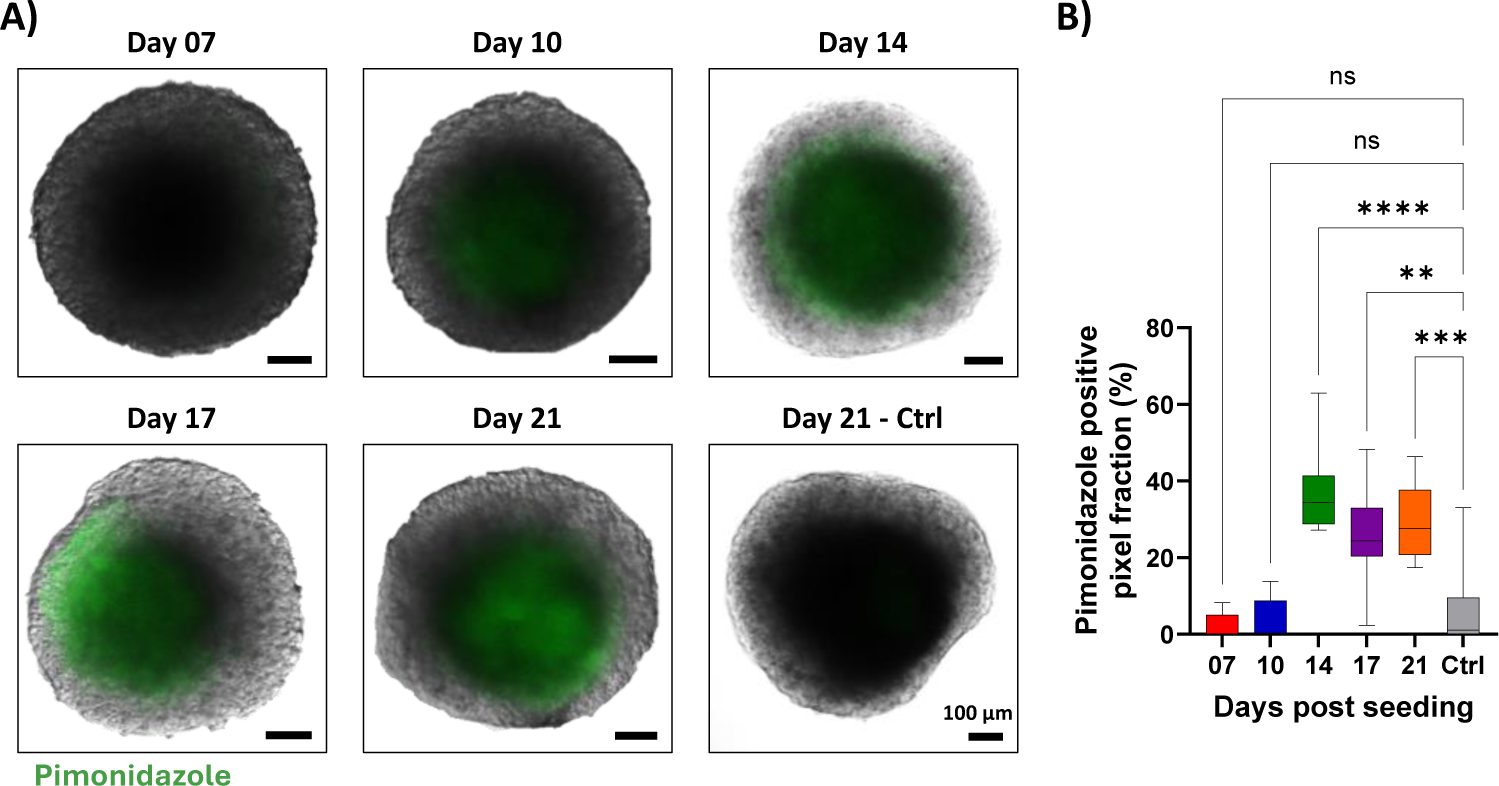
Development of hypoxic regions in FaDu tumor spheroids over time. A) Overlay of brightfield and pimonidazole staining images showing progressive hypoxia from day 7 to 21. (Scale bar = 100 µm). B) Quantification of hypoxic (pimonidazole-positive) area relative to total spheroid area across days (n = 8). Analysis was performed with one-way ANOVA with Dunnet’s multiple comparison test: ns = *p* > 0.05, ** = *p* < 0.01, *** = *p* < 0.001, **** = *p* < 0.0001.

### 4.5. Nanodroplets Alleviates Spheroids Hypoxia

After characterizing the development of hypoxic cores in FaDu spheroids, we sought to evaluate the potential of the PFP NDs to deliver oxygen and overcome this hypoxia. Before performing the oxygen delivery studies, we characterized a commercially available reversible hypoxia fluorescent reagent (Image-iT™ Red Hypoxia Reagent; Thermo Fisher Scientific) for concentration and incubation time. Image-iT™ Red Hypoxia Reagent is a live cell-permeable fluorogenic compound that becomes fluorescent in environments with low oxygen concentrations (< 5%). It is a real-time oxygen detector, and the fluorogenic response reverses with a reversal in oxygen concentrations from hypoxia to normal levels.

Figure 5A shows the untreated and oxygenated PFP NDs treated spheroids before administering the NDs. The spheroids were stained with the hypoxia reagent in red and Hoechst in blue. From the images, we can observe that the core of the spheroids is hypoxic due to the fluorescence of the hypoxia reagent. After the baseline measurement, the oxygenated PFP NDs were added to the treatment group and incubated for 3 hours. After 3 hours, the spheroids were imaged again and are shown in Fig. 5A. While the fluorescence signals of the hypoxia reagent increased in the untreated spheroids, they decreased in the PFP NDs-treated spheroids. This reduction indicates that the PFP NDs diffused into the spheroids, elevating the oxygen levels, and reversing the fluorogenic response of the hypoxia reagent. We further quantified the images for the fluorescence signals before and after the PFP NDs administration and displayed them in Fig. 5B. The quantification revealed a significant decrease in the mean fluorescence intensity of the hypoxia reagent after the PFP NDs treatment compared to untreated controls. After qualitatively showing the hypoxia reduction, we quantified the reduction in the downstream pathway of hypoxia.

**Figure 5:**
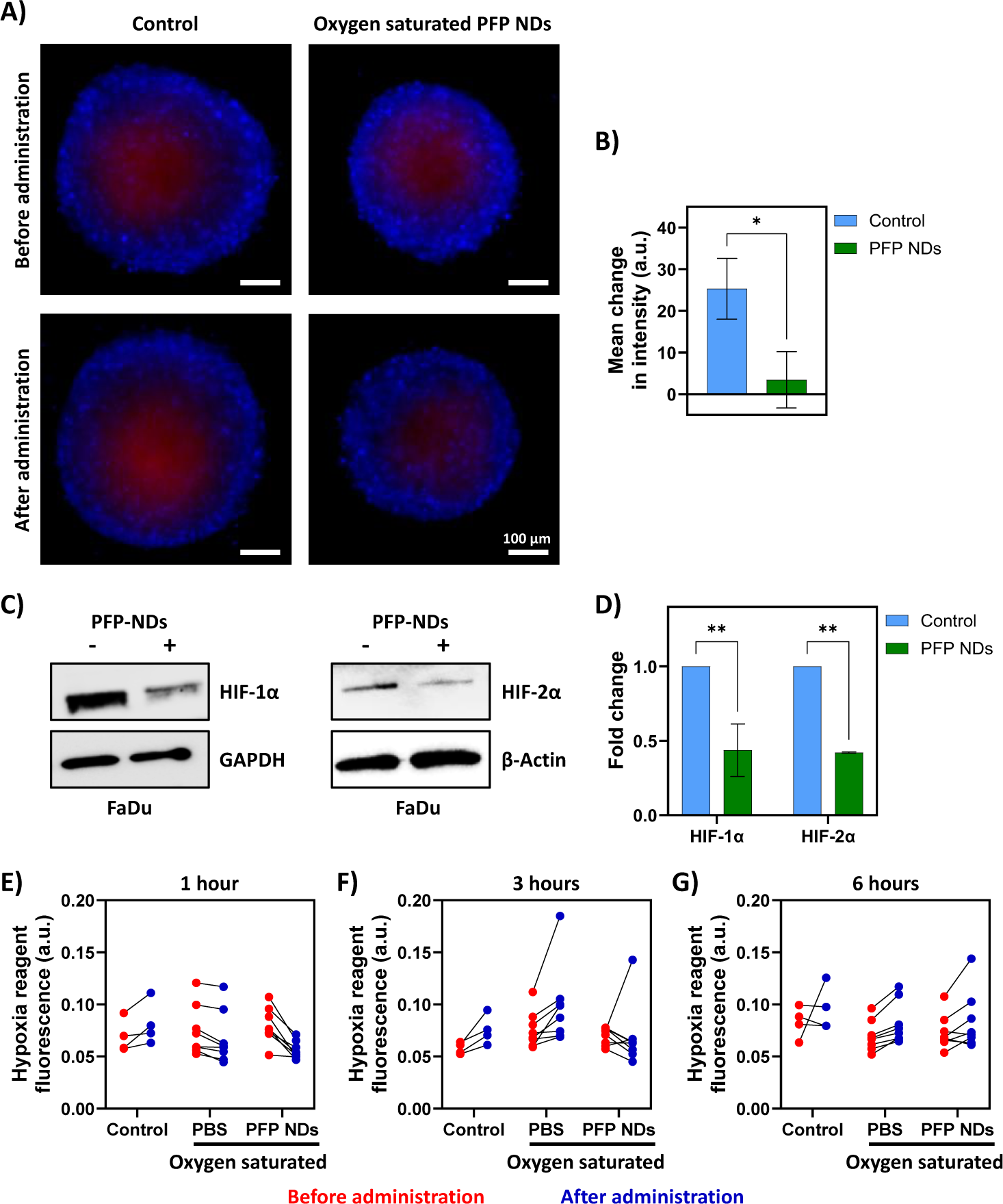
Oxygenated PFP NDs alleviate hypoxia in spheroids. A) Fluorescence images of spheroids stained with Hoechst in blue and hypoxia reagent in red. Control spheroids show increased hypoxia staining, while the spheroid treated with oxygenated PFP NDs for 3 hours show reduced hypoxia staining. (Scale bar = 100 µm). B) Quantification of the mean change in the hypoxia reagent fluorescence intensity, demonstrating reduced hypoxic signal in treated spheroids compared to controls (n = 8). C) Western blot analysis showing decreased levels of HIF-1α and HIF-2α proteins in spheroids treated with oxygenated PFP NDs for 3 hours compared to controls, with GAPDH and β-actin as loading controls. D) Quantification of western blot data, depicting reduced expression of HIF-1α and HIF-2α in treated spheroids compared to controls (n = 3). E-G) Quantification of the hypoxia reagent fluorescence intensity before and after treatment with oxygenated PBS and oxygenated PFP NDs in comparison to untreated control at 1-, 3-, and 6-hours’ time point (n = 4-8). For B), the results are expressed as mean ± standard error. Analysis was performed with unpaired t-test: * = *p* < 0.05. For D), the results are expressed as mean ± standard deviation. Analysis was performed with two-way ANOVA with Šidák’s multiple comparison test: ** = *p* < 0.01.

The hypoxia response pathway is primarily mediated by the HIFs, which are transcription factors that regulate the expression of genes involved in cellular adaptation to low oxygen conditions.^17^ HIFs, primarily HIF-1α and HIF-2α, are regulated by the oxygen-sensing prolyl hydroxylase domain (PHD) enzymes.^77^ Under normoxic conditions, HIFs are hydroxylated by PHDs, leading to their recognition by the von Hippel-Lindau tumor suppressor protein and subsequent proteasomal degradation. However, in hypoxic conditions, the activity of PHDs is inhibited, resulting in the stabilization and accumulation of HIFs.^77^ Consequently, HIF signaling promotes the expression of target genes involved in adaptive responses to hypoxia, such as increased glycolysis, angiogenesis, and cell survival mechanisms.^16,17^ HIFs contribute to the aggressive nature of HNC through various mechanisms, including the promotion of EMT, immune evasion, and the upregulation of drug efflux transporters like P-glycoprotein.^17,78^ Additionally, HIF-mediated metabolic reprogramming towards glycolysis and the induction of angiogenic factors like vascular endothelial growth factor further support tumor growth and metastasis.

To investigate the effects of oxygenated PFP NDs on the hypoxia response pathway, we performed western blot analysis on lysates from spheroids treated with oxygenated PFP NDs for 3 hours and non-treated controls (Fig. 5C). The results revealed a significant downregulation of HIF-1α and HIF-2α expression in the treated spheroids compared to the control group (Fig. 5D). This finding provides compelling evidence for the hypoxia-alleviating effects of the oxygenated PFP NDs. The downregulation of HIFs in response to oxygenated NDs treatment has several implications. Firstly, it suggests that the oxygen delivered by the NDs effectively increases the oxygen levels within the spheroids, leading to the degradation of HIFs under normoxic conditions. This normalization of the hypoxic tumor microenvironment can potentially reduce the aggressive phenotype of HNC cells by suppressing HIF-mediated EMT, immune evasion, and drug efflux transporter upregulation. Secondly, the suppression of the HIF-mediated hypoxic response may sensitize the tumor cells to subsequent therapies hindered by hypoxia, such as PDT or radiotherapy.^32,34,79^ Combining Oxygenated NDs with PDT or radiotherapy could lead to improved therapeutic outcomes for HNC patients.

Finally, we investigated the sustenance of oxygen or in other words the maintenance of oxygenated environments within the core of the spheroids after initial treatment with the oxygenated PFP NDs. We treated the spheroids with hypoxia reagent 24 hours before treatment with oxygenated PBS or PFP NDs for various time points. Figure 5E-G depicts the hypoxia reagent fluorescence of either control spheroids or treated with oxygenated PBS or PFP NDs. Control spheroids, irrespective of the time points, exhibited an increase in hypoxia reagent fluorescence, confirming the continuous presence of hypoxia. The spheroids treated with both oxygenated PBS and PFP NDs at 1-hour time point showed a reduction in the hypoxia reagent fluorescence, indicating successful oxygen delivery. At the 3-hour time point, spheroids treated with oxygenated PBS showed an increase in hypoxia reagent fluorescence, suggesting the consumption of supplied oxygen and the return of hypoxia while majority of the oxygenated PFP NDs treated spheroids (6 out of 8) continued to show a reduction in the fluorescence. This result demonstrates that the oxygen delivered by the NDs can be sustained in the hypoxic tumor spheroids. At the 6-hour time point, all treated groups showed an increase in the fluorescence signals indicating the consumption of the delivered oxygen. These findings demonstrate the ability of the oxygenated PFP NDs not just to alleviate hypoxia but sustain the hypoxia relief for at least 3 hours highlighting their potential to overcome hypoxia-related challenges in HNC treatment.

### 4.6. Optimizing the BPD Delivery to Spheroids by the Nanodroplets

Effective drug delivery to solid tumors remains a significant challenge due to the complex TME, which includes physical barriers such as aberrant vasculature, high interstitial pressure as well as biological barriers like hypoxia, acidity.^17^ Towards the goal of understanding therapeutic outcomes of the nano-drug delivery systems, it is critical to assess *in vitro* drug uptake in complex multi-cellular organoids prior to preclinical assessment. We incubated FaDu and SCC9 spheroids with either free BPD or BPD-PFP NDs and monitored BPD accumulation via fluorescence imaging over time (Fig. 6 A&B) to assess the uptake and penetration of BPD in tumor spheroids. After 1 hour of incubation, both free BPD and BPD-PFP NDs localized primarily in the outer rim region of the spheroids and no significant difference in the uptake was observed. As the incubation time increased, BPD penetration and accumulation into the spheroids’ core gradually improved. Quantification of BPD fluorescence intensity revealed that maximal uptake occurred at 24 hours for both free BPD and BPD-PFP NDs (Fig. C&D). Interestingly, the total BPD accumulated in the spheroids was significantly higher for free BPD compared to BPD-PFP NDs at 3-, 6-, and 24-hours’ time points. The uptake profile of free BPD and BPD-PFP NDs in 3D spheroids were matching the uptake results in 2D monolayer models with free BPD having greater uptake than BPD-PFP NDs (Fig. 2D&E). However, the lower total BPD accumulation observed with BPD-PFP NDs compared to free BPD may be attributed to the slower release kinetics of the NDs formulation. Despite this difference, the ability of PFP NDs to deliver oxygen and BPD to hypoxic tumor microenvironment has been confirmed. This result suggests that a 24-hour incubation period is optimal for maximizing BPD delivery to these tumor spheroids. By combining peak BPD delivery with the supplemental oxygen provided by the PFP NDs, we anticipate enhanced photodynamic therapeutic effects. Further studies are needed to evaluate the therapeutic efficacy of BPD-PFP NDs in hypoxic spheroids and to optimize the treatment protocol for maximum antitumor effects.

**Figure 6:**
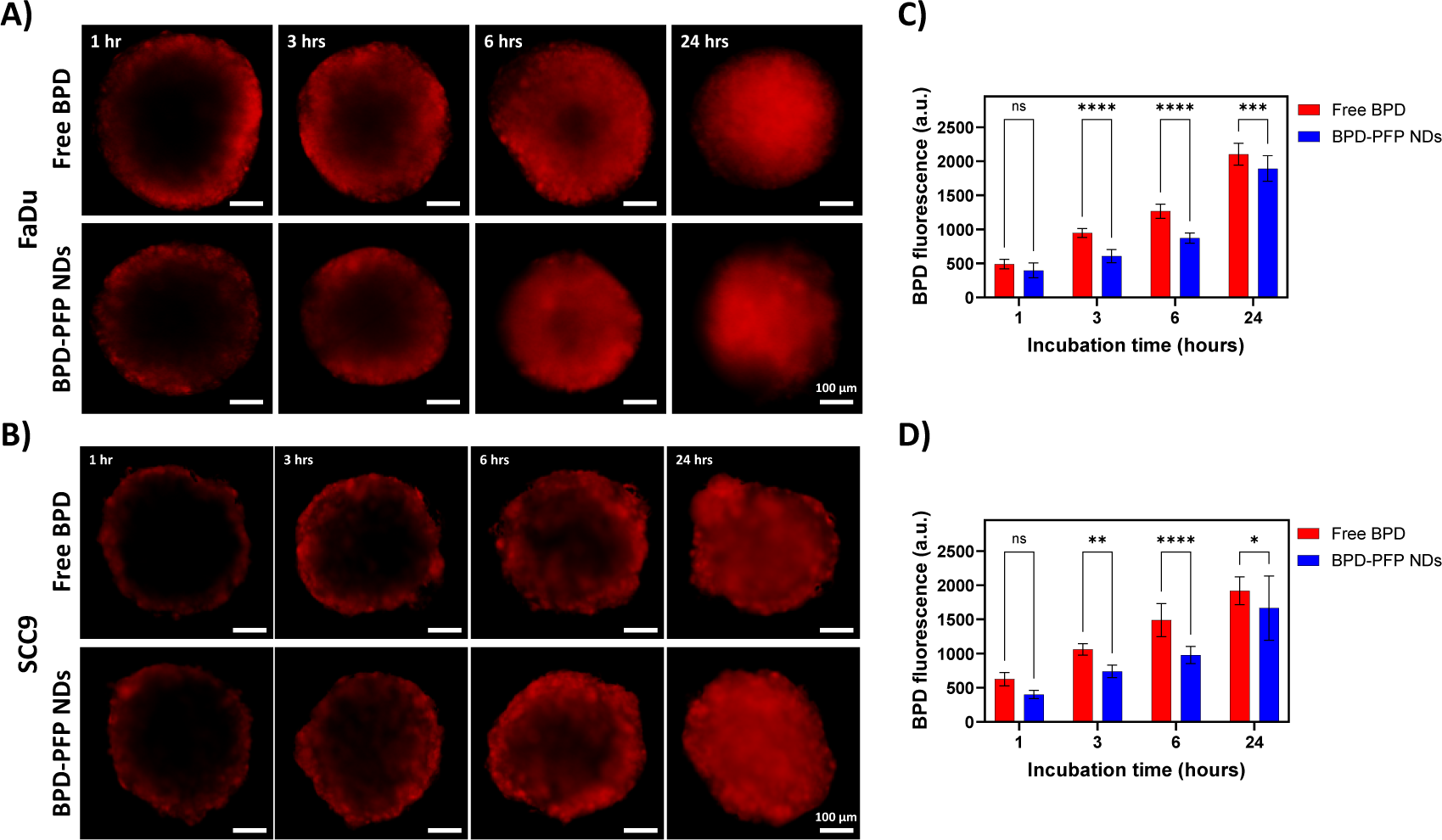
PFP NDs deliver BPD to the spheroids. A & B) The fluorescence images display FaDu or SCC9 spheroids treated with either free BPD or BPD-PFP NDs for 1, 3, 6, and 24 hours. The red fluorescence signal indicates the localization of BPD within the spheroids. (Scale bar = 100 µm). C & D) Quantification of free BPD and BPD-PFP NDs uptake by FaDu or SCC9 spheroids at each time point (n = 16). For C-D), the results are expressed as mean ± standard deviation. Analysis was performed with two-way ANOVA with Šidák’s multiple comparison test: ns = *p* > 0.05, * = *p* < 0.05, ** = *p* < 0.01, *** = *p* < 0.001, **** = *p* < 0.0001.

## 5. Conclusion and Future Work

In this study, we developed and characterized a co-delivery system based on PFP NDs designed to deliver oxygen and therapeutic drugs to hypoxic tumor spheroids. The PFP NDs demonstrated excellent colloidal stability, efficient oxygen loading and release capabilities, and good biocompatibility. Using a 3D multicellular tumor spheroid model, we observed the progression of the hypoxic core over time and validated the ability of oxygenated NDs to alleviate hypoxia within the spheroids. This was evidenced by the reduction in hypoxia reagent fluorescence and the downregulation of hypoxia-inducible factors HIF-1α and HIF-2α. Additionally, we demonstrated the potential of the NDs to deliver drugs effectively to the hypoxic spheroid core. While these results are promising, further *in vitro* and *in vivo* studies are necessary to fully evaluate the therapeutic efficacy of the co-delivery system. Future work will focus on assessing the therapeutic efficacy of PDT when PFP NDs are used to co-deliver oxygen and a PS. Furthermore, exploring the potential of combining this system with other therapeutic modalities could provide a comprehensive treatment strategy for hypoxic HNC tumors.

## Abbreviations

BCA: Bicinchoninic acid
BPD: Benzoporphyrin derivative
DLS: Dynamic light scattering
DMEM: Dulbecco’s Modified Eagle Medium
DMSO: Dimethyl sulfoxide
EMT: Epithelial mesenchymal transition
HIFs: Hypoxia inducible factors
HNC: Head and neck cancer
MTT: (3-(4,5-Dimethylthiazol-2-yl)-2,5-Diphenyltetrazolium Bromide)
NDs: Nanodroplets
PBS: Phosphate buffered saline
PDI: Polydispersity index
PDT: Photodynamic therapy
PEG: Polyethylene glycol
PFC: Perfluorocarbon
PFP: Perfluoropentane
PHD: Prolyl hydroxylase domain
PS: Photosensitizer
TME: Tumor microenvironment
ULA: Ultra-low attachment

## Acknowledgements

The authors would like to gratefully acknowledge funds from the National Institutes of Health R01 CA266701 and Tufts School of Engineering. The authors would also like to acknowledge Dr. Jason Cook for useful discussions on nanodroplets’ synthesis.

## Notes

### Competing Interest Statement

The authors have declared no competing interest.

## References

1 Chow, L. Q. M. Head and Neck Cancer. N Engl J Med 382, 60–72 (2020). 10.1056/NEJMra1715715

2 Bray, F. et al. Global cancer statistics 2018: GLOBOCAN estimates of incidence and mortality worldwide for 36 cancers in 185 countries. CA Cancer J Clin 68, 394–424 (2018). 10.3322/caac.21492

3 Pfister, D. G. et al. Head and Neck Cancers, Version 2.2020, NCCN Clinical Practice Guidelines in Oncology. J Natl Compr Canc Netw 18, 873–898 (2020). 10.6004/jnccn.2020.0031

4 Barsouk, A., Aluru, J. S., Rawla, P., Saginala, K. & Barsouk, A. Epidemiology, Risk Factors, and Prevention of Head and Neck Squamous Cell Carcinoma. Med Sci (Basel) 11 (2023). 10.3390/medsci11020042

5 Ha, P. K., Chang, S. S., Glazer, C. A., Califano, J. A. & Sidransky, D. Molecular techniques and genetic alterations in head and neck cancer. Oral Oncol 45, 335–339 (2009). 10.1016/j.oraloncology.2008.05.015

6 Jawa, Y. et al. Current Insights and Advancements in Head and Neck Cancer: Emerging Biomarkers and Therapeutics with Cues from Single Cell and 3D Model Omics Profiling. Front Oncol 11, 676948 (2021). 10.3389/fonc.2021.676948

7 Whiteside, T. L. Head and Neck Carcinoma Immunotherapy: Facts and Hopes. Clin Cancer Res 24, 6–13 (2018). 10.1158/1078-0432.CCR-17-1261

8 Alsahafi, E. et al. Clinical update on head and neck cancer: molecular biology and ongoing challenges. Cell Death Dis 10, 540 (2019). 10.1038/s41419-019-1769-9

9 Miranda-Galvis, M., Loveless, R., Kowalski, L. P. & Teng, Y. Impacts of Environmental Factors on Head and Neck Cancer Pathogenesis and Progression. Cells 10 (2021). 10.3390/cells10020389

10 Pisani, P. et al. Metastatic disease in head & neck oncology. Acta Otorhinolaryngol Ital 40, S1–S86 (2020). 10.14639/0392-100X-suppl.1-40-2020

11 Hanahan, D. Hallmarks of Cancer: New Dimensions. Cancer Discovery 12, 31–46 (2022). 10.1158/2159-8290.Cd-21-1059

12 Bredell, M. G. et al. Current relevance of hypoxia in head and neck cancer. Oncotarget 7, 50781–50804 (2016). 10.18632/oncotarget.9549

13 Hockel, M. & Vaupel, P. Tumor hypoxia: definitions and current clinical, biologic, and molecular aspects. J Natl Cancer Inst 93, 266–276 (2001). 10.1093/jnci/93.4.266

14 Brown, J. M. & Wilson, W. R. Exploiting tumour hypoxia in cancer treatment. Nat Rev Cancer 4, 437–447 (2004). 10.1038/nrc1367

15 Vaupel, P. & Mayer, A. Hypoxia in cancer: significance and impact on clinical outcome. Cancer Metastasis Rev 26, 225–239 (2007). 10.1007/s10555-007-9055-1

16 Semenza, G. L. Targeting HIF-1 for cancer therapy. Nat Rev Cancer 3, 721–732 (2003). 10.1038/nrc1187

17 Jing, X. et al. Role of hypoxia in cancer therapy by regulating the tumor microenvironment. Mol Cancer 18, 157 (2019). 10.1186/s12943-019-1089-9

18 Melissaridou, S. et al. The effect of 2D and 3D cell cultures on treatment response, EMT profile and stem cell features in head and neck cancer. Cancer Cell Int 19, 16 (2019). 10.1186/s12935-019-0733-1

19 Shi, R., Liao, C. & Zhang, Q. Hypoxia-Driven Effects in Cancer: Characterization, Mechanisms, and Therapeutic Implications. Cells 10 (2021). 10.3390/cells10030678

20 Harada, H. Hypoxia-inducible factor 1-mediated characteristic features of cancer cells for tumor radioresistance. J Radiat Res 57 **Suppl 1**, i99–i105 (2016). 10.1093/jrr/rrw012

21 Celli, J. P. et al. Imaging and photodynamic therapy: mechanisms, monitoring, and optimization. Chem Rev 110, 2795–2838 (2010). 10.1021/cr900300p

22 Graham, K. & Unger, E. Overcoming tumor hypoxia as a barrier to radiotherapy, chemotherapy and immunotherapy in cancer treatment. Int J Nanomedicine 13, 6049–6058 (2018). 10.2147/IJN.S140462

23 Ahn, P. H. et al. Lesion oxygenation associates with clinical outcomes in premalignant and early stage head and neck tumors treated on a phase 1 trial of photodynamic therapy. Photodiagnosis Photodyn Ther 21, 28–35 (2018). 10.1016/j.pdpdt.2017.10.015

24 Larue, L. et al. Fighting Hypoxia to Improve PDT. Pharmaceuticals (Basel) 12 (2019). 10.3390/ph12040163

25 Zhao, L. et al. Advanced nanotechnology for hypoxia-associated antitumor therapy. Nanoscale 12, 2855–2874 (2020). 10.1039/c9nr09071a

26 Xu, M. et al. Smart strategies to overcome tumor hypoxia toward the enhancement of cancer therapy. Nanoscale 12, 21519–21533 (2020). 10.1039/d0nr05501h

27 Shen, Z. et al. Strategies to improve photodynamic therapy efficacy by relieving the tumor hypoxia environment. NPG Asia Materials 13 (2021). 10.1038/s41427-021-00303-1

28 Du, Y., Han, J., Jin, F. & Du, Y. Recent Strategies to Address Hypoxic Tumor Environments in Photodynamic Therapy. Pharmaceutics 14 (2022). 10.3390/pharmaceutics14091763

29 Mohanto, N., Park, Y. J. & Jee, J. P. Current perspectives of artificial oxygen carriers as red blood cell substitutes: a review of old to cutting-edge technologies using in vitro and in vivo assessments. J Pharm Investig 53, 153–190 (2023). 10.1007/s40005-022-00590-y

30 Dias, A. M. A., Freire, M., Coutinho, J. A. P. & Marrucho, I. M. Solubility of oxygen in liquid perfluorocarbons. Fluid Phase Equilibria 222-223, 325-330 (2004). 10.1016/j.fluid.2004.06.037

31 Kakaei, N., Amirian, R., Azadi, M., Mohammadi, G. & Izadi, Z. Perfluorocarbons: A perspective of theranostic applications and challenges. Frontiers in Bioengineering and Biotechnology 11 (2023). 10.3389/fbioe.2023.1115254

32 Xavierselvan, M. et al. Photoacoustic nanodroplets for oxygen enhanced photodynamic therapy of cancer. Photoacoustics 25, 100306 (2022). 10.1016/j.pacs.2021.100306

33 Sheng, D., Deng, L., Li, P., Wang, Z. & Zhang, Q. Perfluorocarbon Nanodroplets with Deep Tumor Penetration and Controlled Drug Delivery for Ultrasound/Fluorescence Imaging Guided Breast Cancer Therapy. ACS Biomater Sci Eng 7, 605–616 (2021). 10.1021/acsbiomaterials.0c01333

34 Cheng, Y. et al. Perfluorocarbon nanoparticles enhance reactive oxygen levels and tumour growth inhibition in photodynamic therapy. Nat Commun 6, 8785 (2015). 10.1038/ncomms9785

35 Hu, H. et al. Perfluorocarbon-based O(2) nanocarrier for efficient photodynamic therapy. J Mater Chem B 7, 1116–1123 (2019). 10.1039/c8tb01844h

36 Hu, D. et al. Perfluorocarbon-Loaded and Redox-Activatable Photosensitizing Agent with Oxygen Supply for Enhancement of Fluorescence/Photoacoustic Imaging Guided Tumor Photodynamic Therapy. Advanced Functional Materials 29 (2019). 10.1002/adfm.201806199

37 Wilhelm, E. & Battino, R. Thermodynamic functions of the solubilities of gases in liquids at 25.deg. Chemical Reviews 73, 1–9 (1973). 10.1021/cr60281a001

38 Geyer, M. in talic>Praxis · Theorie · Variationen · Leitungstechnik · Forschung · Entwicklung und Anwendung in verschiedenen Ländern Berufspolitik · Kritische Glosse (eds Jürgen Körner, Herbert Neubig, & Ulrich Rosin) 31-35 (Springer Berlin Heidelberg, 1988).

39 Wang, Z. et al. Oxygen-Delivering Polyfluorocarbon Nanovehicles Improve Tumor Oxygenation and Potentiate Photodynamic-Mediated Antitumor Immunity. ACS Nano 15, 5405–5419 (2021). 10.1021/acsnano.1c00033

40 Jagers, J., Wrobeln, A. & Ferenz, K. B. Perfluorocarbon-based oxygen carriers: from physics to physiology. Pflugers Arch 473, 139–150 (2021). 10.1007/s00424-020-02482-2

41 Riess, J. G. Understanding the fundamentals of perfluorocarbons and perfluorocarbon emulsions relevant to in vivo oxygen delivery. Artif Cells Blood Substit Immobil Biotechnol 33, 47–63 (2005). 10.1081/bio-200046659

42 Zhao, A., Lee, J. & Emelianov, S. Formulation and Acoustic Modulation of Optically Vaporized Perfluorocarbon Nanodroplets. J Vis Exp (2021). 10.3791/62814

43 Hannah, A., Luke, G., Wilson, K., Homan, K. & Emelianov, S. Indocyanine green-loaded photoacoustic nanodroplets: dual contrast nanoconstructs for enhanced photoacoustic and ultrasound imaging. ACS Nano 8, 250–259 (2014). 10.1021/nn403527r

44 Loskutova, K., Grishenkov, D. & Ghorbani, M. Review on Acoustic Droplet Vaporization in Ultrasound Diagnostics and Therapeutics. Biomed Res Int 2019, 9480193 (2019). 10.1155/2019/9480193

45 Wang, J. et al. All-in-One Theranostic Nanoplatform Based on Hollow MoS(x) for Photothermally-maneuvered Oxygen Self-enriched Photodynamic Therapy. Theranostics 8, 955–971 (2018). 10.7150/thno.22325

46 Kapalczynska, M. et al. 2D and 3D cell cultures - a comparison of different types of cancer cell cultures. Arch Med Sci 14, 910–919 (2018). 10.5114/aoms.2016.63743

47 Lv, D., Hu, Z., Lu, L., Lu, H. & Xu, X. Three-dimensional cell culture: A powerful tool in tumor research and drug discovery. Oncol Lett 14, 6999–7010 (2017). 10.3892/ol.2017.7134

48 Yamada, K. M. & Cukierman, E. Modeling tissue morphogenesis and cancer in 3D. Cell 130, 601–610 (2007). 10.1016/j.cell.2007.08.006

49 Aguilar Cosme, J. R., Gagui, D. C., Bryant, H. E. & Claeyssens, F. Morphological Response in Cancer Spheroids for Screening Photodynamic Therapy Parameters. Front Mol Biosci 8, 784962 (2021). 10.3389/fmolb.2021.784962

50 Hirschhaeuser, F. et al. Multicellular tumor spheroids: an underestimated tool is catching up again. J Biotechnol 148, 3–15 (2010). 10.1016/j.jbiotec.2010.01.012

51 Desai, N. Challenges in Development of Nanoparticle-Based Therapeutics. The AAPS Journal 14, 282–295 (2012). 10.1208/s12248-012-9339-4

52 Phan, H. T. & Haes, A. J. What Does Nanoparticle Stability Mean? The Journal of Physical Chemistry C 123, 16495–16507 (2019). 10.1021/acs.jpcc.9b00913

53 Liu, Y., Zhu, S., Gu, Z., Chen, C. & Zhao, Y. Toxicity of manufactured nanomaterials. Particuology 69, 31–48 (2022). 10.1016/j.partic.2021.11.007

54 Lea-Banks, H. et al. Ultrasound-sensitive nanodroplets achieve targeted neuromodulation. Journal of Controlled Release 332, 30–39 (2021). 10.1016/j.jconrel.2021.02.010

55 Song, R., Peng, C., Xu, X., Zou, R. & Yao, S. Facile fabrication of uniform nanoscale perfluorocarbon droplets as ultrasound contrast agents. Microfluidics and Nanofluidics 23, 12 (2019). 10.1007/s10404-018-2172-z

56 Chibowski, E. & Szcześ, A. Zeta potential and surface charge of DPPC and DOPC liposomes in the presence of PLC enzyme. Adsorption 22, 755–765 (2016). 10.1007/s10450-016-9767-z

57 Neunert, G., Tomaszewska-Gras, J., Witkowski, S. & Polewski, K. Tocopheryl Succinate-Induced Structural Changes in DPPC Liposomes: DSC and ANS Fluorescence Studies. Molecules 25 (2020).

58 Obaid, G., Jin, W., Bano, S., Kessel, D. & Hasan, T. Nanolipid Formulations of Benzoporphyrin Derivative: Exploring the Dependence of Nanoconstruct Photophysics and Photochemistry on Their Therapeutic Index in Ovarian Cancer Cells. Photochem Photobiol 95, 364–377 (2019). 10.1111/php.13002

59 Bhandari, C. et al. PD-L1 Immune Checkpoint Targeted Photoactivable Liposomes (iTPALs) Prime the Stroma of Pancreatic Tumors and Promote Self-Delivery. Advanced Healthcare Materials **n/a**, 2304340 (2024). 10.1002/adhm.202304340

60 Bruun, K. & Hille, C. Study on intracellular delivery of liposome encapsulated quantum dots using advanced fluorescence microscopy. Scientific Reports 9, 10504 (2019). 10.1038/s41598-019-46732-5

61 Tunuguntla, R. et al. Lipid Bilayer Composition Can Influence the Orientation of Proteorhodopsin in Artificial Membranes. Biophysical Journal 105, 1388–1396 (2013). 10.1016/j.bpj.2013.07.043

62 Huang, H. C. et al. Photodynamic Therapy Synergizes with Irinotecan to Overcome Compensatory Mechanisms and Improve Treatment Outcomes in Pancreatic Cancer. Cancer Res 76, 1066–1077 (2016). 10.1158/0008-5472.CAN-15-0391

63 Lea-Banks, H., Wu, S.-K., Lee, H. & Hynynen, K. Ultrasound-triggered oxygen-loaded nanodroplets enhance and monitor cerebral damage from sonodynamic therapy. Nanotheranostics 6, 376–387 (2022). 10.7150/ntno.71946

64 Owen, J. et al. Orally administered oxygen nanobubbles enhance tumor response to sonodynamic therapy. Nano Select 3, 394–401 (2022). 10.1002/nano.202100038

65 Chen, C.-Y. & Chen, C.-Y. Targeted and Oxygen-Enriched Nanoplatform for Enhanced Photodynamic Therapy: In Vitro 2D Cell and 3D Spheroid Model Evaluation. Macromolecular Bioscience 23, 2300196 (2023). 10.1002/mabi.202300196

66 Lambert, E. & Janjic, J. M. Quality by design approach identifies critical parameters driving oxygen delivery performance in vitro for perfluorocarbon based artificial oxygen carriers. Scientific Reports 11, 5569 (2021). 10.1038/s41598-021-84076-1

67 Petrovic, L. Z. et al. Mutual impact of clinically translatable near-infrared dyes on photoacoustic image contrast and in vitro photodynamic therapy efficacy. J Biomed Opt 25, 1–12 (2020). 10.1117/1.JBO.25.6.063808

68 Hafiz, S. S. et al. Eutectic Gallium–Indium Nanoparticles for Photodynamic Therapy of Pancreatic Cancer. ACS Applied Nano Materials 5, 6125–6139 (2022). 10.1021/acsanm.1c04353

69 Meng, D. et al. Dual-sensitive and highly biocompatible O-carboxymethyl chitosan nanodroplets for prostate tumor ultrasonic imaging and treatment. Cancer Nanotechnology 14, 19 (2023). 10.1186/s12645-023-00172-z

70 Zhou, X. et al. Ultrasound-responsive highly biocompatible nanodroplets loaded with doxorubicin for tumor imaging and treatment in vivo. Drug Delivery 27, 469–481 (2020). 10.1080/10717544.2020.1739170

71 Immordino, M. L., Dosio F Fau - Cattel, L. & Cattel, L. Stealth liposomes: review of the basic science, rationale, and clinical applications, existing and potential.

72 Pinto, B., Henriques, A. C., Silva, P. M. A. & Bousbaa, H. Three-Dimensional Spheroids as In Vitro Preclinical Models for Cancer Research. Pharmaceutics 12 (2020).

73 Close, D. A. & Johnston, P. A. Detection and impact of hypoxic regions in multicellular tumor spheroid cultures formed by head and neck squamous cell carcinoma cells lines. SLAS Discov 27, 39–54 (2022). 10.1016/j.slasd.2021.10.008

74 Riffle, S., Pandey, R. N., Albert, M. & Hegde, R. S. Linking hypoxia, DNA damage and proliferation in multicellular tumor spheroids. BMC Cancer 17, 338 (2017). 10.1186/s12885-017-3319-0

75 Däster, S. et al. Induction of hypoxia and necrosis in multicellular tumor spheroids is associated with resistance to chemotherapy treatment. Oncotarget (2017). 10.18632%2Foncotarget.13857

76 Grimes, D. R., Kelly, C., Bloch, K. & Partridge, M. A method for estimating the oxygen consumption rate in multicellular tumour spheroids. Journal of The Royal Society Interface 11, 20131124 (2014). 10.1098/rsif.2013.1124

77 Masoud, G. N. & Li, W. HIF-1alpha pathway: role, regulation and intervention for cancer therapy. Acta Pharm Sin B 5, 378–389 (2015). 10.1016/j.apsb.2015.05.007

78 Robinson, K. & Tiriveedhi, V. Perplexing Role of P-Glycoprotein in Tumor Microenvironment. Front Oncol 10, 265 (2020). 10.3389/fonc.2020.00265

79 Eisenbrey, J. R. et al. Sensitization of Hypoxic Tumors to Radiation Therapy Using Ultrasound-Sensitive Oxygen Microbubbles. International Journal of Radiation Oncology*Biology*Physics 101, 88–96 (2018). 10.1016/j.ijrobp.2018.01.042

